# Assessment of the optimum fertilizer rates and planting density for soybean production in China

**DOI:** 10.1101/688986

**Authors:** Jilong Lv, Ping He, Dan Wei, Xinpeng Xu, Shaojun Qiu, Shicheng Zhao

## Abstract

Fertilization rate and planting density are important factors affecting crop yield. A large number of soybean [*Glycine max* (L.) Merr] field experimental data (1998-2017) were collected through different database sources to evaluate the optimum fertilizer rate and planting density for high yield of spring and summer soybean in China. The yield of spring and summer soybean gradually increased over year, with their average yields were 2610 and 2724 kg ha^−1^, respectively. Based on the fitted quadratic curve, the optimal rate of nitrogen (N), phosphorus (P), and potassium (K) fertilizers for high yield of summer soybean was 96 kg N ha^−1^, 80 kg P_2_O_5_ ha^−1^, and 126 kg K_2_O ha^−1^, and the corresponding yields were 3038, 2801 and 2305 kg ha^−1^, respectively. The optimal rate of N, P and K fertilizers for spring soybean was 71 kg N ha^-1^, 108 kg P_2_O_5_ ha^-1^ and 74 kg K_2_O ha^−1^, and the corresponding yields were 2932, 2834 and 2678 kg ha^−1^, respectively. The optimum density was 27×10^4^ and 34×10^4^ plants ha^−1^ under high yield for summer and spring soybean, respectively. Stepwise regression analysis showed that the P fertilizer had the greatest influence on the spring soybean yield followed by K fertilizer and planting density. For summer soybean, population density had the major effect on yield followed by P fertilizer. Overall, the P fertilization and planting density should be payed attention to increase soybean yield in different regions of China.

## Introduction

Soybean [*Glycine max* (L.) Merr] is an important source of high-quality plant protein and edible oil. With its biological nitrogen-fixing (BNF) capacity, soybean is an important crop in decreasing N fertilizer application and sustaining high crop yield in the crop rotation system [1]. Soy-based foods such as tofu, soy milk, soy sauce, and miso have been developed for human consumption [2], and soybean meal is used as animal feed. As an oil crop, soybean also has a wide range of uses in the industrial field in addition to its home usage [3].

The current total soybean yield is approximately 363 million tons globally, with an average yield of 2782 kg ha^−1^[4], but still cannot meet the need of growing population. The factors influencing high soybean yield include climatic conditions, soil characteristics, soybean varieties, nutrient management and cultivation practices [5]. Different climatic conditions (such as temperature variation, sunshine duration, and precipitation) affect the planting date, grain-filling period and total growth period of crop that ultimately influence the crop yield [6,7]. Soil characteristics can determine soil nutrient supply capacity and affect crop growth and yield, because the 2/3 of nutrients absorbed during the crop growing period came from soil and only 1/3 of nutrients supplied by fertilizers [8]. Nitrogen (N), phosphorus (P), and potassium (K) are primarily essential nutrients for crop growth [9–11], and balanced fertilization play an important role in increasing crop yield [12]. The soybean seed with high protein contents has a large demand for N. The appropriate amount of N fertilizer can enhance the photosynthesis, improve the seed yield, protein, and oil per unit area. P is involved in the formation of new cells structures, protein synthesis, transportation and transformation of organic compounds in soybean [13]. P fertilization can effectively improve nodular symbiosis between legumes and rhizobium species [14,15], with enhancement of the activity of nitrogenase enzyme during nodulation, and results in N fixation ability of root nodules [16]. K is mainly involved in the formation of carbohydrates in soybean and increases the strength of stems, and its deficiency prolongs the maturation period and decreases the quality and yield [17]. Appropriate planting density is essential for increasing soybean yield, because it improves the utilization of light energy in leaves, promotes the nutrient absorption, and increases the dry matter accumulation with yield [18].

China is an important soybean producer and consumer country in the world, but its soybean production is far from meeting the demand of increasing population, with a deficit of 8-10× 10^7^ tons that imports from other countries every year [19]. In China, soybean cultivation is divided into spring and summer soybean according to their sowing time. The spring soybean is mainly concentrated in northeast China with a monoculture, and the summer soybean is mainly in the north-central China and south China plains [20], with mainly wheat-soybean rotation. Because of low seed yield (average 1773 kg ha^−1^) [20], farmers pay less attention to soybean production in China even knowing its importance. At the same time, it is believed that soybean can fix N through root nodules and does not need to apply ample amount of N fertilizer; however, the BNF cannot meet the entire N demand of soybean, especially under high yield. A large number of field studies on optimum soybean fertilization and planting density were carried out with some recommendations on fertilization rate and planting density to improve crop yield and nutrient use efficiency [21,22]. However, these experiments were conducted in individual fields and recommendations only reflect the individual fields for optimal nutrient management or density. This individual test result is not universal for large regions, because there are great variations in climate and soil conditions across soybean production areas of China, and it is necessary to conclude the universal management measure in fertilization and planting density by summing up earlier study datasets. Therefore, we collected a large number of experimental data including fertilization rate, planting density, and seed yield of soybean across different production regions from 1993 to 2017. The objective is to obtain the optimum N, P, and K fertilization rate and planting density with consideration of higher yield in main soybean planting areas of China.

## Materials and methods

### Data sources

The database used in this study included field experiments conducted by the International Plant Nutrition Institute (IPNI) China Program, the Soybean Industrial System Research Database, and papers published in the China Knowledge Network (CNKI) journal from 1998 to 2017. The field studies did not involve endangered or protected species, so no specific permissions were required for the location/activity. The keyword used to access the literature included: soybean, yield, density, and fertilization rate. Total 748 field experimental data were analyzed in this study, these experiments included “3414” balanced fertilization experiment, population density and fertilizer rate experiments (Table1). The distribution of experimental sites was showed in Fig.1. All field trials were included clear fertilization rates, plant population density, and seed yield. The soybean test varieties were widely grown locally.

**Fig 1.**
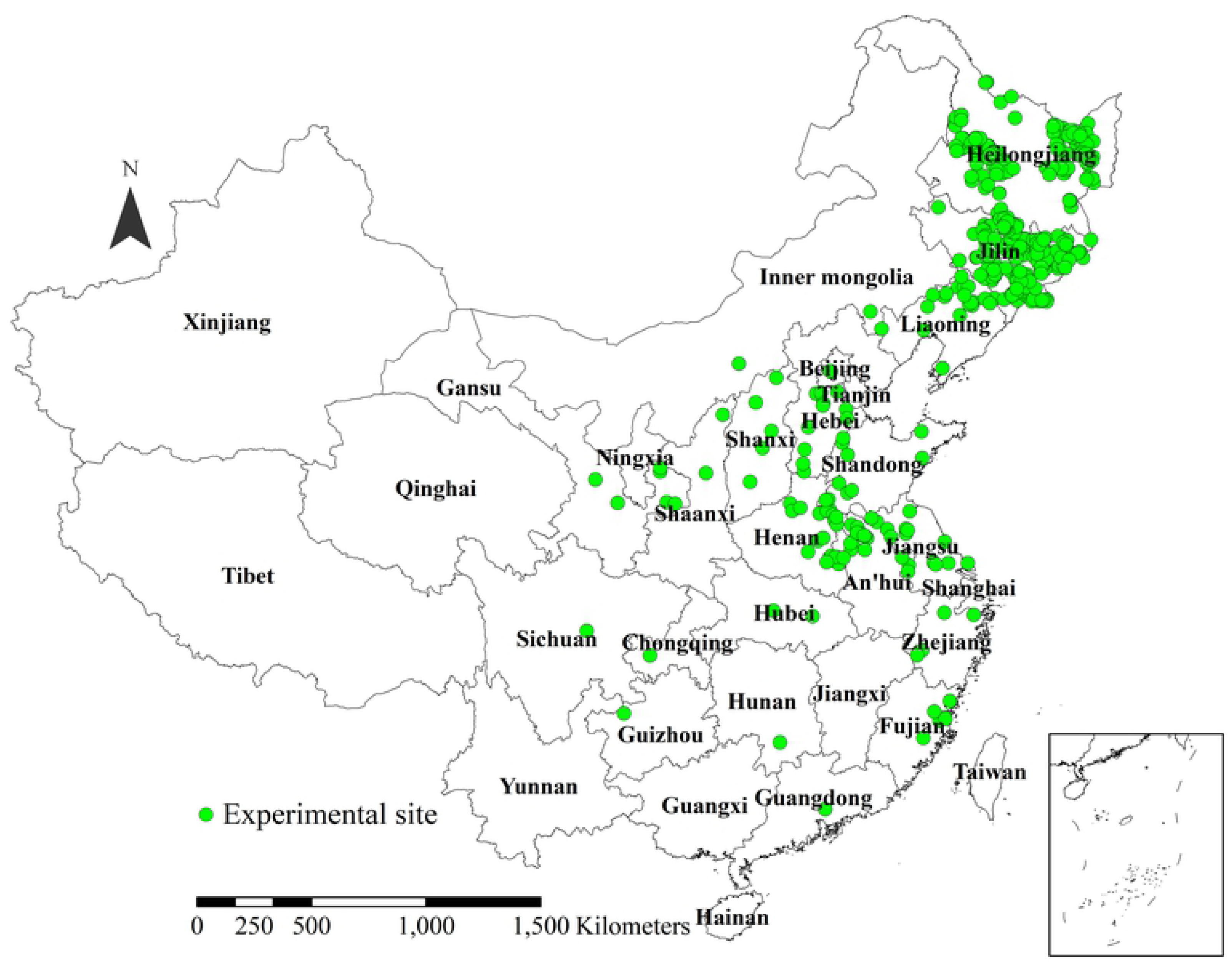
The location distribution of experiment sites in China.

**Table1.** The type and number of experiment for soybean data collection.

| Experiment types | Number of experiment |
| --- | --- |
| “3414” and balanced fertilization | 635 |
| Density and fertilization rate | 24 |
| Nitrogen fertilizer rate | 27 |
| Phosphorus fertilizer rate | 20 |
| Potassium fertilizer rate | 29 |
| Density experiment | 13 |

### Data Analysis

Soybean data was divided into summer and spring soybean based on the sowing date (summer soybean is sown in early June and spring soybean is sown in later April or early May) for further analysis. The data with harvest index value less than 0.4 kg kg^−1^ were excluded because they were assumed that the crop suffered abiotic or biotic stresses other than nutrient deficiency during the growing season [1]. The soybean seed yield derived from the optimum practical treatment (N, P, and K were recommended based on soil testing) was used to evaluate the yield change and average yield within 20 years. Soybean seed yield was adjusted to 135 g kg^−1^ moisture content. The quadratic function of seed yield corresponding to different rates of N, P, K fertilizers, and planting density were fitted to determine the optimum rate for fertilization and planting density with Microsoft Excel. Stepwise multiple regression analysis (*p* < 0.05) was applied to detect the factors influencing yield by using SPSS 19.0 version (SPSS, Inc., Chicago, IL, USA).

## Results

### Soybean yield change

The seed yield of spring and summer soybean gradually increased from 1998 to 2017, and the increased rate was higher in summer than spring soybean, and the summer soybean presented higher average seed yield (2724 kg ha^−1^) relative to spring soybean (2610 kg ha^−1^) (Fig.2). The yield of summer and spring soybean were mainly concentrated ranged in 2000–3000 kg ha^−1^, presenting 46.4% and 59.8% of the yield data followed by 3,000-4,000 kg ha^−1^, accounting for 31.4% and 23.1%, respectively (Fig. 3). The yield variation of summer soybean was larger than that of spring soybean.

**Fig 2.**
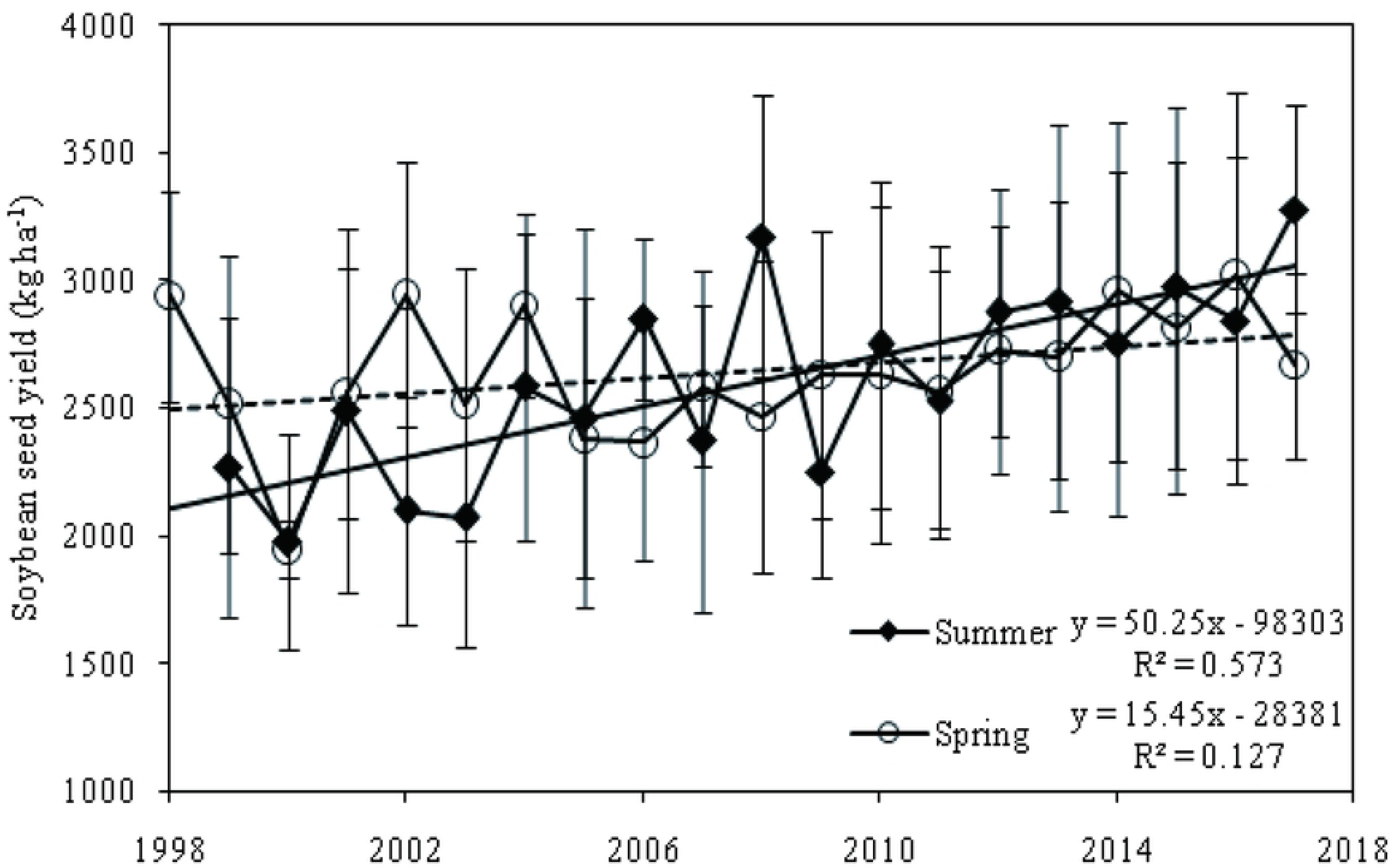
Change in average seed yield for summer and spring soybean from 1998 to 2017. (Yield date from optimum fertilization treatment. Trend line: solid line for summer soybean and dotted line for spring soybean). Means ±standard deviation (*n* = 3) are shown.

**Fig 3.**
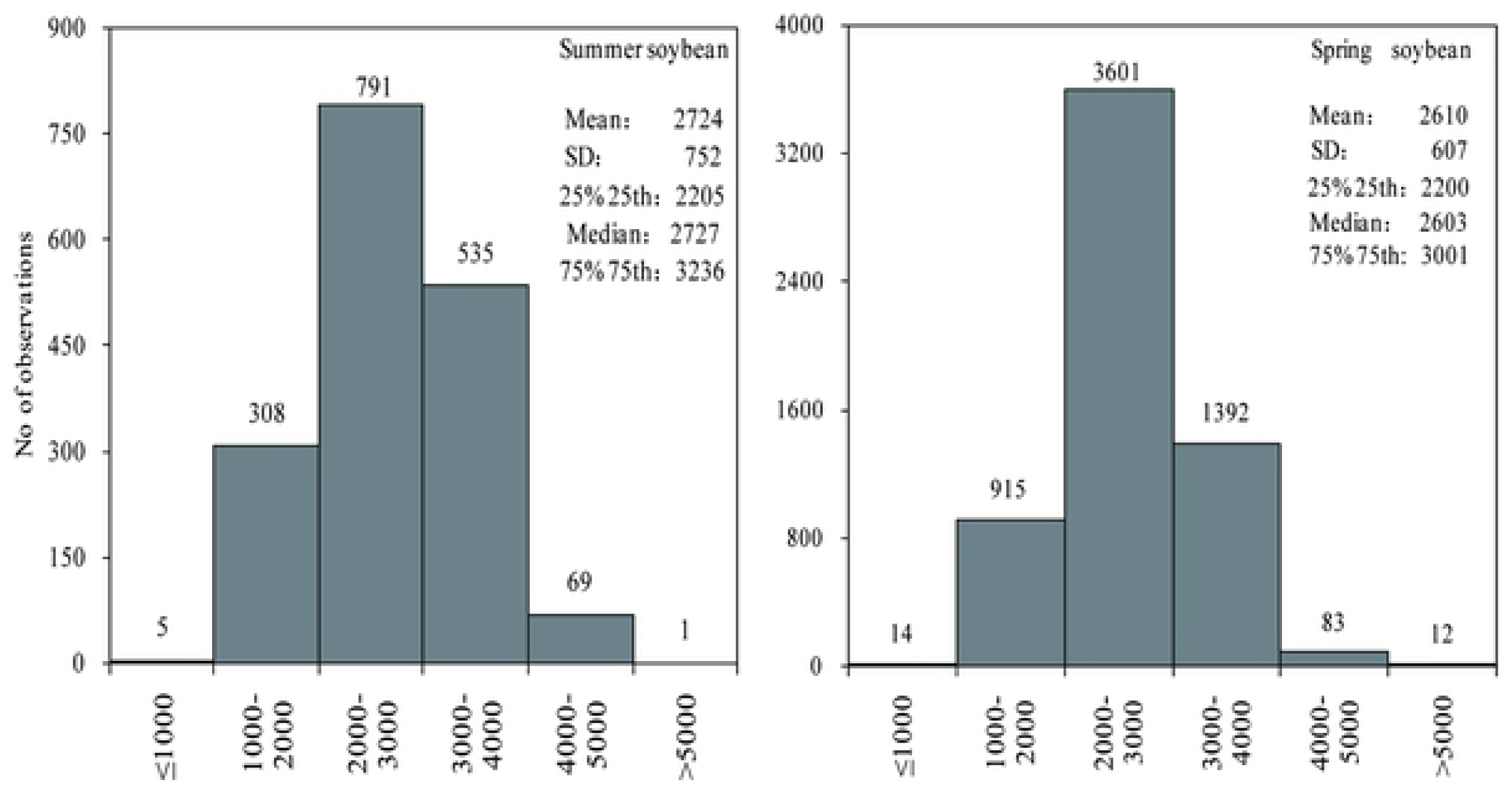
Sequences distribution of soybean yield from 1998 to 2017 (Yield date from optimum fertilization treatment)

#### Relationship between soybean yield and fertilization rates

Soybean seed yield showed an increasing-decreasing change with the increasing rate of N, P, and K fertilizers (Fig. 4). According to the quadratic equation fitted between the fertilizer rate and soybean yield, when the maximum yield of summer soybean was obtained, the rate of N, P, and K fertilizers was 96 kg N ha^−1^, 80 kg P_2_O_5_ ha^−1^, and 126 kg K_2_O ha^−1^, respectively, with the corresponding maximum yield 3038, 2801 and 2305 kg ha^−1^, respectively. For the spring soybean, the rate of N, P, and K fertilizers was 71 kg N ha^−1^,108 kg P_2_O_5_ ha^−1^, 74 kg K_2_O ha^−1^, respectively, and the corresponding maximum yield was 2932, 2834 and 2678 kg ha^−1^, respectively.

**Fig 4.**
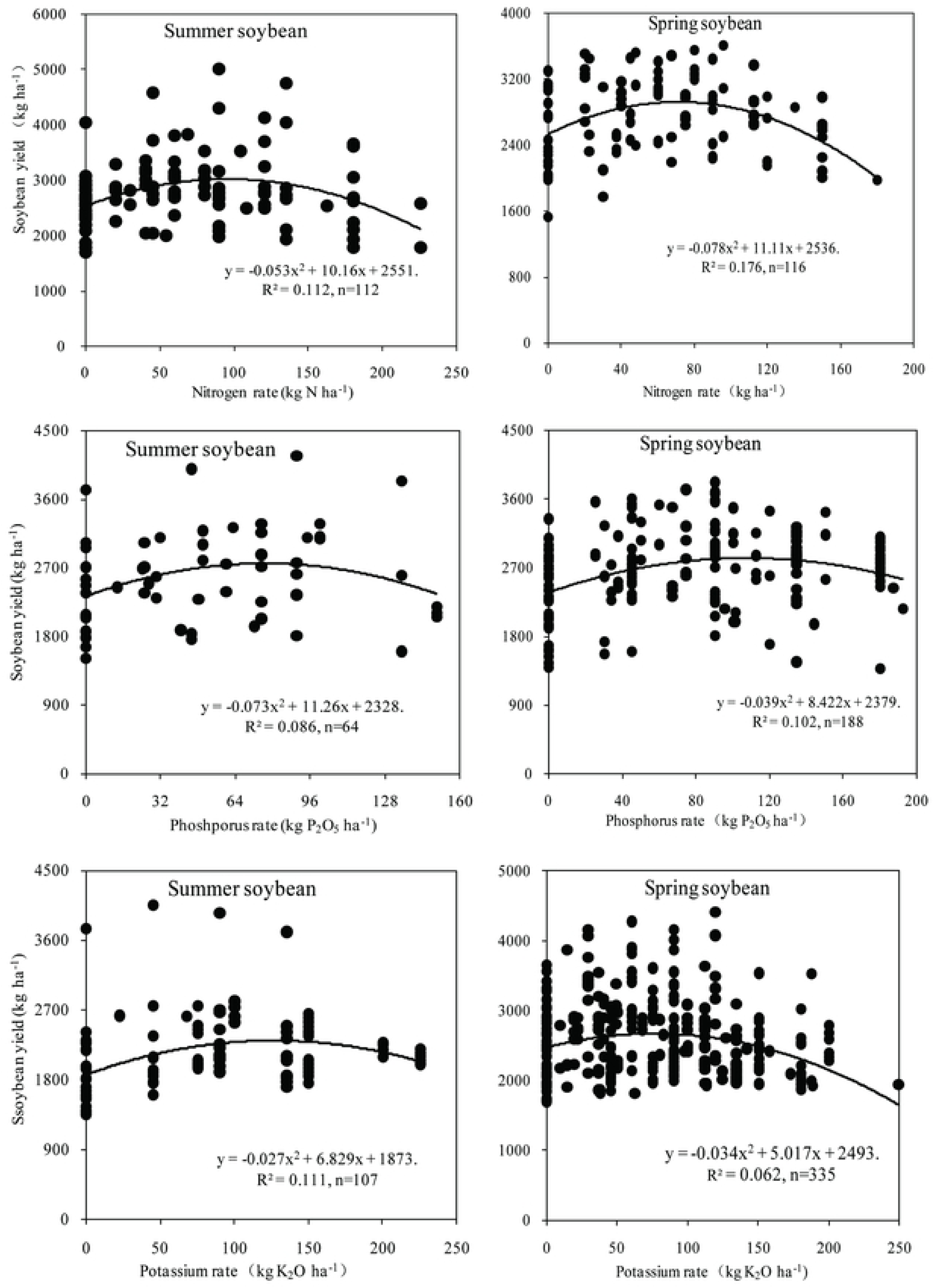
Relationship between soybean seed yield and the rate of N, P, and K fertilization (1998-2017).

### Relationship between planting density and seed yield

The soybean yield also showed an increasing-decreasing change with the increase in planting density (Fig. 5). According to the fitted curve, the yield of summer and spring soybean was the highest under the density of 26×10^4^ and 34×10^4^ plants ha^−1^, the corresponding yields were 2936 and 2791 kg ha^−1^, respectively. The summer soybean presented higher rate in increase or decrease of yield relative to spring soybean with increasing planting density, indicating that the yield of summer soybean was greatly affected by the change in population density relative to spring soybean.

**Fig 5.**
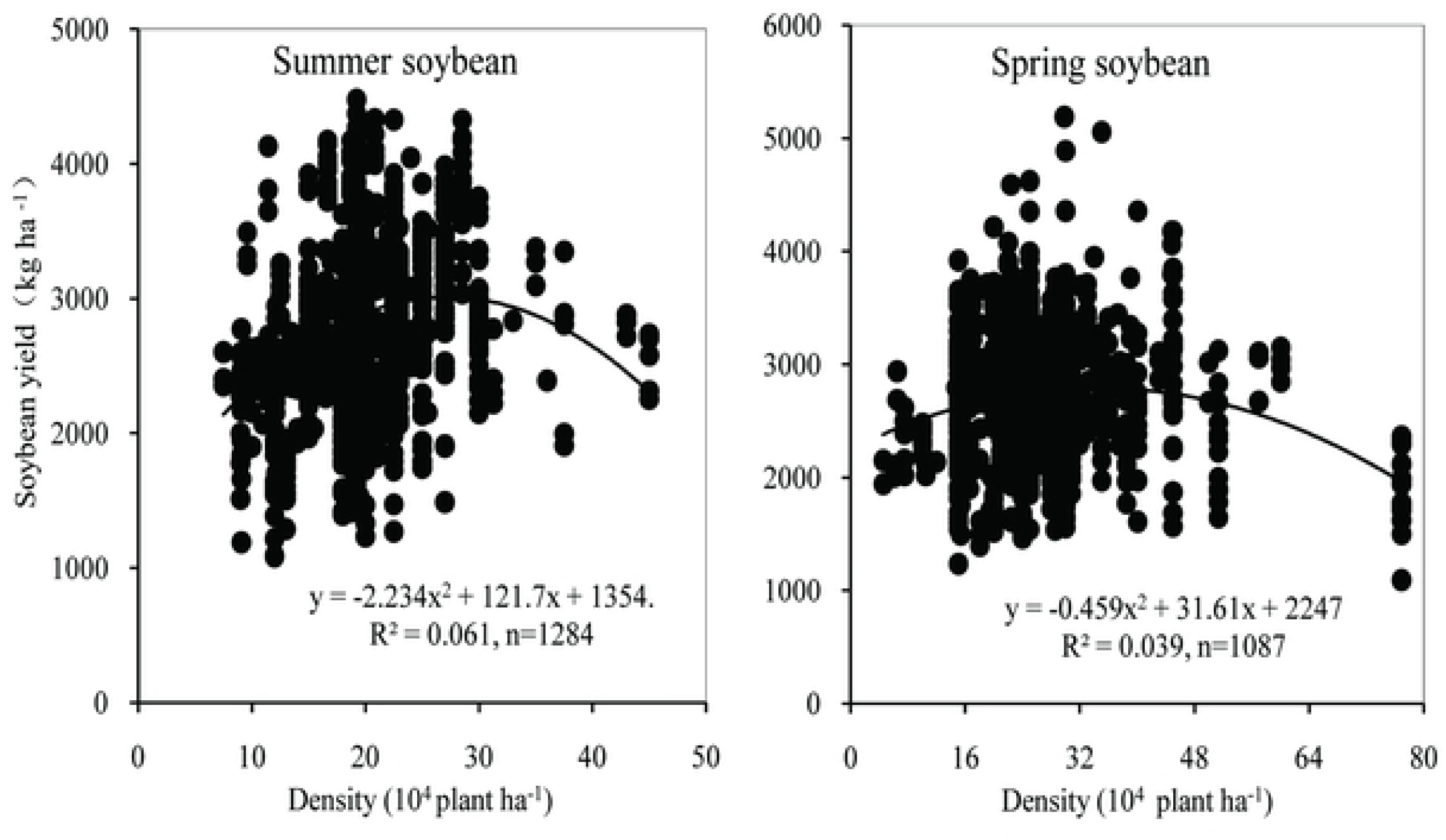
Relationship between soybean yield and planting density.

### Relationship between fertilization rate, planting density, and soybean yield

Stepwise regression analysis showed that density had the greatest effect on summer soybean yield, followed by the P fertilizer rate (Table 2). For spring soybean, the P fertilizer had the greatest effect on yield, followed by K, N fertilizers and planting density. However, the introduction of density in the regression curve indicated that planting density and P fertilizer played important roles in high yield, followed by K fertilizer in spring soybean area.

**Table 2.** Stepwise regression analysis of soybean yield (Y) and planting density, the rate of N, P, and K fertilizers (X).

| Dependent | independents | Summer soybean |  | Spring soybean |  |
| --- | --- | --- | --- | --- | --- |
|  |  | R value | P value | R value | P value |
| Soybean<br>yield<br>(Y) | Density (X1) | 0.578 | p=0 | 0.267 | p=0.021 |
|  | N rate (X2) | 0.061 | p=0.454 | 0.331 | p=0.001 |
|  | P rate (X3) | 0.325 | p=0 | 0.426 | p=0 |
|  | K rate (X4) | 0.041 | p=0.169 | 0.343 | p=0 |
| Regression equation |  | Y=2071.92+28.68X1+1.92X3,<br>R= 0.661 |  | Y=2341.63+11.03X1-3.86X3+6.08X4,<br>R= 0.526 |  |

## Discussion

Due to the genetic improvement in soybean variety, the application of balanced fertilization and other agricultural technologies since 1998, the soybean yield increased over years in China. The average yield of summer soybean was higher relative to spring soybean, which was mainly related to regional soil fertility and climatic condition. Soybean is a short-day thermophilic crop and is sensitive to light and temperature [23–25]. Spring soybean growed in northeast China, which has low average temperature and low rainfall in soybean growing season. The summer soybean region has high temperature and precipitation, which is more conducive to the soybean growth. Meanwhile, the fertilization rate and soil nutrient content in farmland of north-central and south China are higher than those in northeast China [26]. The average yield data might be overestimated in this study, because these data are from the experimental fields in the main soybean producing areas. The fertilization rate, planting density, and other management practices were scientific in these experimental fields; and in fact, the insufficient and unbalanced fertilization and the low planting density extensive existed in farmers’ fields of these soybean production areas.

Rational application of N, P and K nutrients is the key factor attaining high yield [27–29]. This study showed that the optimal rate of N, P, K for summer soybean was 96 kg N ha^−1^, 80 kg P_2_O_5_ ha^−1^, 126 kg K_2_O ha^−1^, and these values were 71 kg N ha^-1^, 108 P_2_O_5_ ha^−1^, and 74 kg K_2_O ha^−1^ for spring soybean, respectively. These data are deviated from previous results, probably because the earlier researchers have proposed the appropriate amount of fertilizer based on the results of a single test, only for specific areas. Our study summarized the multi-year and multi-point test data in different regions, the number of fertilization treatment and sample were large, the fertilization gradient is dense and with strong reliability. The optimum N and K fertilizer rates of summer soybean were higher than that of spring soybean. Firstly, the higher attainable yield of summer soybean required more nutrients; secondly, the soil N a d K contents were higher in north-central relative to northeast China [30]. Soil P are surplus in most area of China, and the surplus was greater in north-central than northeast China [30–32], while the low temperature in early spring season limits soil P availability, which cannot meet the P demand of soybean in early growth stag. In order to meet the P demand for high soybean yield, people increased the P fertilizer inputs; leading to the calculated optimum P fertilizer rate in spring soybean was higher than that in summer soybean [33–34]. Therefore, how to activate and make full use of accumulated soil P in early spring in northeast China is also an important research direction for reducing P fertilization and increasing P use efficiency.

The optimum density of spring soybean is higher than that of summer soybean, because the optimum density of crops is determined by varieties, local climate and environmental conditions, etc. Soybean close-planting can improve the interception and utilization of light and increase soybean yield. The spring temperature and the annual effective accumulated temperature are low in northeast China, it is necessary to increase planting density to make full use of light and heat to attain high yield.

Regression analysis showed that P fertilizer and density had great effects on the high yield of spring and summer soybeans. Because P is an important element for the synthesis of protein and fat in soybean seed, and P can promote the formation and development of root nodules, the fixation of atmospheric N, the conversion of ammonia and the formation of amino acids [35,36], as well as promote the absorption of N and K in soybean [37]. In addition, low soil P availability resulted from low temperature is also an important reason in spring soybean region. In north-central China, the most of the soils is calcareous with high K content, and straw return is prevalent and can effectively supplement soil K supply; however, in northeast China, few of crop straw were returned to soils because of its slow degradation under low temperature, soil K was rapidly consumed due to crop uptake. Appropriate density can increase the photosynthetic rate and nutrient absorption of soybean per unit area, thus increasing soybean yield [38]. In north-central China, soybean was sowed directly with any tillage after wheat harvest, a large number of wheat straw returned to the fields affects the quality of soybean sowing and seedling rate, the soybean density is general low in actual production [39].

The summer and spring soybean yields under optimum N, P and K rates were higher than the average soybean yield except for the yield under optimum K rate for summer soybean. At the same time, the summer and spring soybean yields under the optimum density were higher than their average yields. The results showed that optimum fertilizer application and reasonable planting density increased soybean yield. Therefore, we should make optimum fertilization rates and planting density based on different plant areas to increase yield in the future soybean planting. However, due to the large variation in soil texture and fertility in spring and summer soybean planting areas, a constant fertilization rate and planting density may not be suitable across the whole areas, we can adjust these values based on the idea of “large formula and small adjustment” [40–42] combining with the characters of different regions to attain high soybean yield.

## Conclusions

Our study found that the seed yield of spring and summer soybean increased over year since 1998 in China, with a higher average yield for summer soybean as compared to spring soybean. The optimum rate of N, P and K fertilizers under high yield was 96 kg N ha^−1^, 80 kg P_2_O_5_ ha^−1^ and 126 kg K_2_O ha^−1^ for summer soybean, and was 71 kg N ha^−1^, 108 kg P_2_O_5_ ha^−1^, 74 kg K_2_O ha^−1^ for spring soybean, respectively. The optimum population density for high yield of summer soybean was higher than that of spring soybean. Planting density is a key factor for high yield of soybean and need to be increased to attain high yield in both soybean producing areas. We should pay attention to P fertilization in both soybean areas, and the attention also is needed for K fertilization in the spring soybean area.

## Acknowledgements

This project was supported by the National Key Research & Development Program of China (2018YFD0201001 and 2016YFD0200102).

